# Network expansion of genetic associations defines a pleiotropy map of human cell biology

**DOI:** 10.1101/2021.07.19.452924

**Authors:** Inigo Barrio-Hernandez, Jeremy Schwartzentruber, Anjali Shrivastava, Noemi del-Toro, Qian Zhang, Glyn Bradley, Henning Hermjakob, Sandra Orchard, Ian Dunham, Carl A. Anderson, Pablo Porras, Pedro Beltrao

## Abstract

Proteins that interact within molecular networks tend to have similar functions and when perturbed influence the same organismal traits. Interaction networks can be used to expand the list of likely trait associated genes from genome-wide association studies (GWAS). Here, we used improvements in SNP-to-gene mapping to perform network based expansion of trait associated genes for 1,002 human traits showing that this recovers known disease genes or drug targets. The similarity of network expansion scores identifies groups of traits likely to share a common genetic basis as well as the biological processes underlying this. We identified 73 pleiotropic gene modules linked to multiple traits that are enriched in genes involved in processes such as protein ubiquitination and RNA processing. We show examples of modules linked to human diseases enriched in genes with pathogenic variants found in patients or relevant mouse knock-out phenotypes and can be used to map targets of approved drugs for repurposing opportunities. Finally, we illustrate the use of the network expansion scores to study genes at inflammatory bowel disease (IBD) GWAS loci, and implicate IBD-relevant genes with strong functional and genetic support.

## Introduction

Proteins that interact tend to take part in the same cellular functions and be important for the same organismal traits (Oti and Brunner 2007; Carter, Hofree, and Ideker 2013). Through a principle of guilt-by-association, it has been shown that molecular networks can be used to predict the function or disease relevance of human genes (Oti et al. 2006; Franke et al. 2006; Vanunu et al. 2010). Based on this, physical or functional interaction networks can augment genome-wide association studies (GWAS) by using GWAS-linked genes as seeds in a network to identify additional trait-associated genes (H. Fang et al. 2019; Lee et al. 2011; Greene et al. 2015; Huang et al. 2018). It is well known that GWAS loci are enriched in genes encoding for successful drug targets (Nelson et al. 2015; Mountjoy et al. 2020). While genes linked to a trait by network expansion are not necessarily within GWAS linked loci, these are also enriched for successful drug targets even when excluding the genes with direct genetic support (MacNamara et al. 2020).

This an opportune time to revisit the application of network approaches to GWAS interpretation, based on recent large improvements in: the human molecular networks available; the approaches for SNP to gene mapping; and the extent of human traits/diseases mapped by GWAS. In particular, there have been substantial improvements in the identification of likely causal genes within GWAS loci using expression and protein quantitative trait loci analysis (Zhu et al. 2016; Sun et al. 2018), as well as machine learning based integrative approaches (Mountjoy et al. 2020).

The genetic study of large numbers of diverse human traits also opens the door for the study of pleiotropy, which occurs when a single genetic change affects multiple traits. Studying pleiotropy can help in the drug discovery process to either increase the number of potential indications for a drug or to avoid unwanted side-effects. Yeast studies of pleiotropy, based on gene deletion, have revealed pleiotropic cellular processes that include endocytosis, ubiquitin system, stress response and protein folding, amino acid biosynthesis, and global transcriptional regulation, among others (Hillenmeyer et al. 2008). Human GWAS data have been extensively used to quantify pleiotropy at SNP level for different traits (Boyle, Li, and Pritchard 2017; Watanabe et al. 2019; Hackinger and Zeggini 2017). While this has shed light into the degree of pleiotropy and the relation between traits this has not often led to the identification of the biological processes and mechanisms that underlie their common genetic basis.

Here we have used recent advances in SNP-to-gene mapping and a comprehensive protein interaction network to augment GWAS data for 1,002 traits by network expansion. This network expansion recovers known disease genes not associated by GWAS, it identifies groups of traits under the influence of the same cellular processes and defines a pleiotropy map of human cell biology. We show examples of gene modules linked to human diseases enriched for genetic variants found in patients and used to map drug targets for possible repurposing opportunities. Finally, we illustrate the use of the network expansion scores to characterize inflammatory bowel disease (IBD) genes at GWAS loci, and implicate IBD-relevant genes with strong functional and genetic support.

## Results

### Systematic augmentation of GWAS with network propagation

We aimed to improve the identification of trait-associated genes and processes via network propagation of GWAS information. Recent studies have shown that a comprehensive protein interaction network is critical for network propagation efforts (Huang et al. 2018). Here, we combined the IMEx physical protein interaction dataset (Porras et al. 2020) from IntAct (protein-protein interactions) (Orchard et al. 2014), Reactome (pathways) (Jassal et al. 2020) and Signor (directed signalling pathways) (Licata et al. 2020). To facilitate the re-use of this physical interaction data we have made it available via a Neo4j Graph Database that can be queried to extract different sub-components including subsetting by the source of interaction (e.g. Reactome, Signor) or type of interaction (e.g. directed, signed) (ftp://ftp.ebi.ac.uk/pub/databases/intact/various/ot_graphdb/current). The physical interactions were further combined with high confidence functional associations from the STRING database (v11)(Szklarczyk et al. 2019) to a final combined network containing 571,917 edges connecting 18,410 total proteins (nodes) (**Fig 1A**).

**Figure 1.**
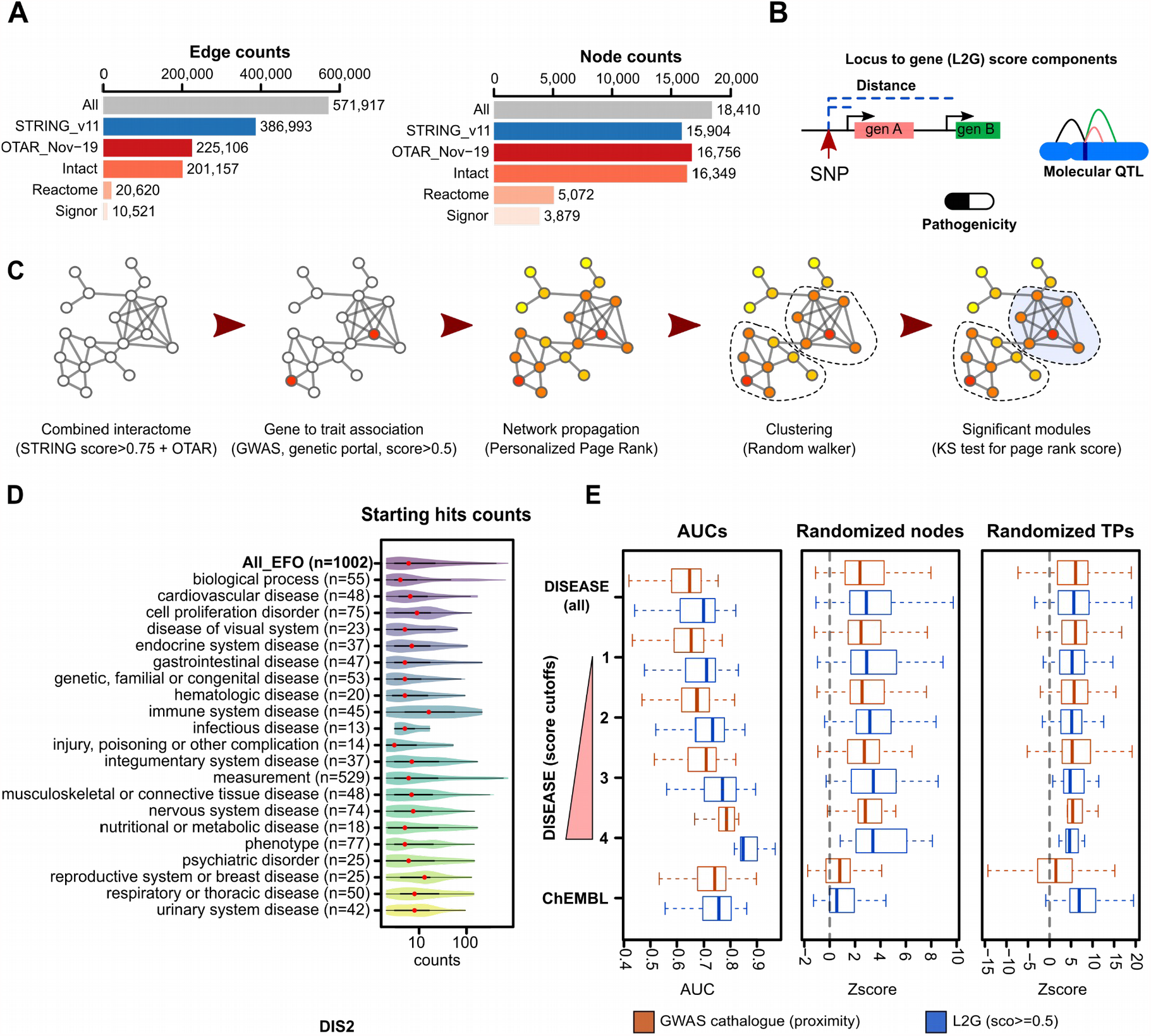
Implementation and benchmarking of network based augmentation of GWAS. A) Edge and node counts of the combined interactome and its components B) Graphic representation of some Locus-to-gene score (L2G) components: SNP to gene distance, data from QTLs, and variant effect predictions. C) Graphical representation of the network-based approach: network propagation of the initial input, clustering using a random walker to find gene communities, and scoring of modules using the distribution of page rank score D) Number of starting genes linked to traits, grouped in therapeutic areas. E) Benchmarking of the method, using as a starting signal all genes from the GWAS catalog (red boxplots) and the genes from the Open Targets genetics portal with L2G score bigger than 0.5. The Area under the ROC curves (AUCs) are calculated using as positive hits DISEASE database, with increasing cut-off values for its gene to trait score (see methods), as well as clinical trials data from CHEMBL (clinical phase II or higher). We also re-calculated the AUCs and determined Z-scores after node randomization of the network keeping the same degree as well as true positives.

The simplest approach to link a SNP reported in a GWAS to a likely causal gene is to select the closest gene. However, the closest gene may not be the causal gene and the integration of expression quantitative trait loci (eQTL) and other data has proven to be useful for SNP to gene mapping (de Lange et al. 2017). Here we mapped GWAS trait associations to genes using the Locus-to-gene (L2G) score from Open Targets Genetics, a recently developed machine learning approach that integrates SNP fine-mapping, gene distance, and molecular QTL information to identify causal genes (**Fig 1B**) (Mountjoy et al. 2020). Genes with L2G scores higher than 0.5 are expected to be causal for the respective trait association in 50% of cases.

For each GWAS, we used genes with L2G > 0.5 as seed genes for the interaction network. Of 7,660 GWAS genes linked to at least one trait, 7248 correspond to proteins present in the interaction network. We then used the Personalized Page Rank (PPR) algorithm to score all other protein coding genes represented in the interaction network. Genes connected via short paths to GWAS genes receive higher network propagation scores (**Fig 1C**). Genes in the top 25% of network propagation scores were used to identify gene modules (see Methods), from which we selected those significantly enriched for high network propagation scores (BH adjusted *p-value*<0.05 with Kolmogorov–Smirnov test) and with at least 2 GWAS linked genes (see Methods). We applied this approach to 1,002 traits (see list in **STable 1**) with GWAS in the Open Targets Genetics portal that had at least 2 genes mapped to the interactome. These GWAS were spread across 21 therapeutic areas, and differed in the number of GWAS-linked genes (median 6, range 2-763) (**Fig 1D**).

In order to measure the capacity of the network expansion to recover trait associated genes, we defined a “gold standard” set of genes known to be associated with human diseases (from diseases.jensenlab.org) or which are known drug targets for specific human diseases (from ChEMBL, see Methods). For the disease associated genes, we further stratified these based on confidence levels (see Methods). To avoid circularity in benchmarking the network expansion approach, we excluded gold standard genes that overlapped with GWAS-linked genes for the respective diseases. The network propagation score predicted disease-associated genes for both gold standard gene sets with an average area under the receiver operating curve (ROC) greater than 0.7 for the most stringent definition of disease-associated genes as well as known drug targets (**Fig 1E**). For comparison we also seeded the network with genes linked to a trait by proximity to the associated SNP (GWAS catalog). Overall, performance was modestly better when using SNP to gene mappings that integrate across diverse data than when using only gene proximity to the lead SNP (**Fig 1E**). This is consistent with the observation that SNP to gene distance is one the strongest predictors of causal genes (Stacey et al. 2019). The observed capacity to identify known disease genes or drug targets was significantly higher than observed with random permutation of gene names in the network or in the gold standard gene sets (**Fig 1E**, node and TP permutations). This suggests the observed performance is not strongly biased by the placement of the gold standard genes within the network.

Overall we obtained network propagation scores for 1,002 traits and gene modules for 906 traits (**STable 1**). In the next sections we illustrate the usefulness of these for the study of genetics of human traits and diseases.

### Network propagation identifies human traits influenced by the same biological processes

Identifying groups of traits that are likely to have a common genetic and biological basis is of value because drugs used to treat one disease may be relevant for other related diseases. Genetic sharing between human traits is often determined from genome-wide genetic correlation of summary statistics from GWAS; however, this approach does not identify how the shared genetics corresponds to shared biological processes. In addition many GWAS do not report the full summary statistics needed for such comparisons. In contrast, network propagation scores can be calculated from results available for all GWAS and be used to identify traits influenced by the same biological processes. To benchmark trait-trait associations derived from network propagation, we used annotations for human traits captured in the Experimental Factor Ontology (EFO). For example, pairs of related neurological traits will tend to share a higher than average number of annotation terms in EFO. Using these annotations we defined 796 pairs of traits that are functionally related and therefore likely to have a common genetic and biological basis (see Methods). Using this benchmark we can show that the similarity in the network propagation scores can identify functionally related pairs of traits (**SFig 1**).

To explore trait-trait relationships based on the similarity of their perturbed biological processes, we used the pairwise distance of network propagation scores to build a tree by hierarchical clustering (**Fig 2A**), and defined 54 sub-groups of traits. The traits tend to group according to functional similarity with 34 out of 54 having an EFO term annotated to over 50% of the traits in the group (**Fig 2A**). We illustrate in **Fig 2B** examples of traits that are grouped together according to the network propagation scores. These include known relationships between immune associated traits such as cellulitis or psoriasis and immunoglobulin measurements (IgG); the relationship between skin neoplasms and skin pigmentation or eye colour; or the clustering of cardiovascular diseases (acute coronary symptoms) with lipoproteins measurements and cholesterol. The latter group links together plasminogen levels in plasma with aortic stenosis and peripheral vascular system conditions.

**Figure 2.**
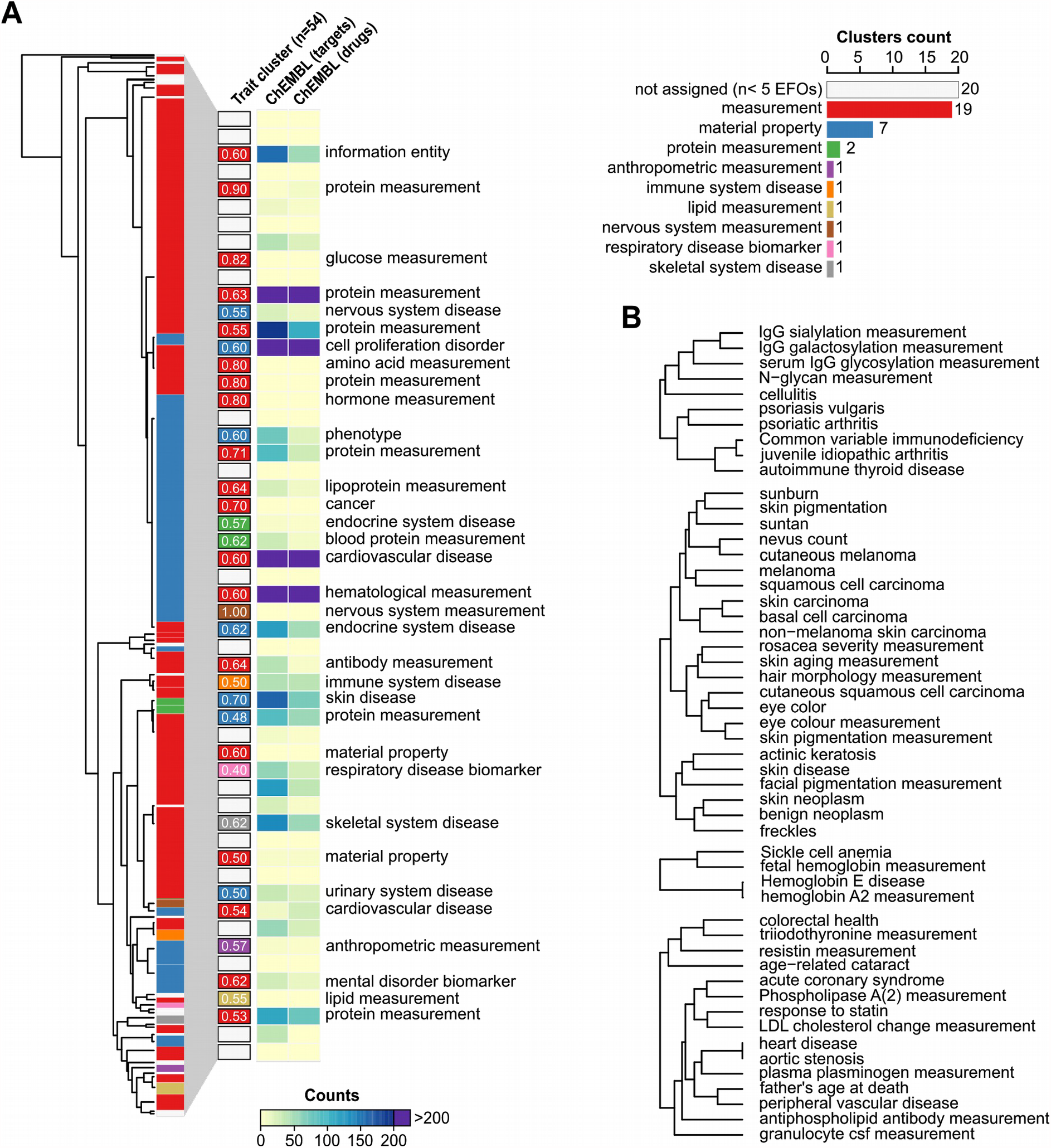
Trait-trait genetic and functional similarities determined from network expansion of GWAS data. A) Tree showing the Manhattan distance between all traits, using the full PPR score. Hierarchical clustering was performed using h=1 cut off, leading to 54 clusters, coloured depending on the predominant EFO ancestry term. In the right panel, barplot showing the 54 clusters with the frequencies for the predominant EFO ancestry terms and a heatmap showing the counts for ChEMBL targets and drugs. B) Examples of traits grouped together using the Manhattan distance, extracted from the tree in panel A.

We obtained drug indications from the ChEMBL database for the diseases in each cluster (**Fig 2A**). This allows us to find clusters where drugs may be considered for repurposing as well as groups of traits where drug development is most needed. 18 clusters representing 64 traits contain no associated drug and represent less well explored areas of drug development. These trait clusters, genes and corresponding drugs are available in **STable 1**. In addition, as we show in the next section, we can use the network propagation to identify the biological processes whose perturbation underlie the trait-trai similarities.

### Pleiotropy of gene modules across human traits

We can study the pleiotropy of human cell biology by identifying which of the above described gene modules tend to be associated with many human traits. This allows us to understand how perturbations in specific aspects of cell biology may have broad consequences across multiple traits. In total we found 2021 associations between gene modules and traits, from which 886 (43.8%) are gene modules linked to a single trait and the remaining can be collapsed to 73 gene modules linked to 2 or more traits (**Fig 3A, STable 2**, see Methods). The modules associated with more than one trait did not have a significantly larger number of genes compared to those linked to single traits (*p-value*= 0.72, kolmogorov Smirnov test).

**Figure 3.**
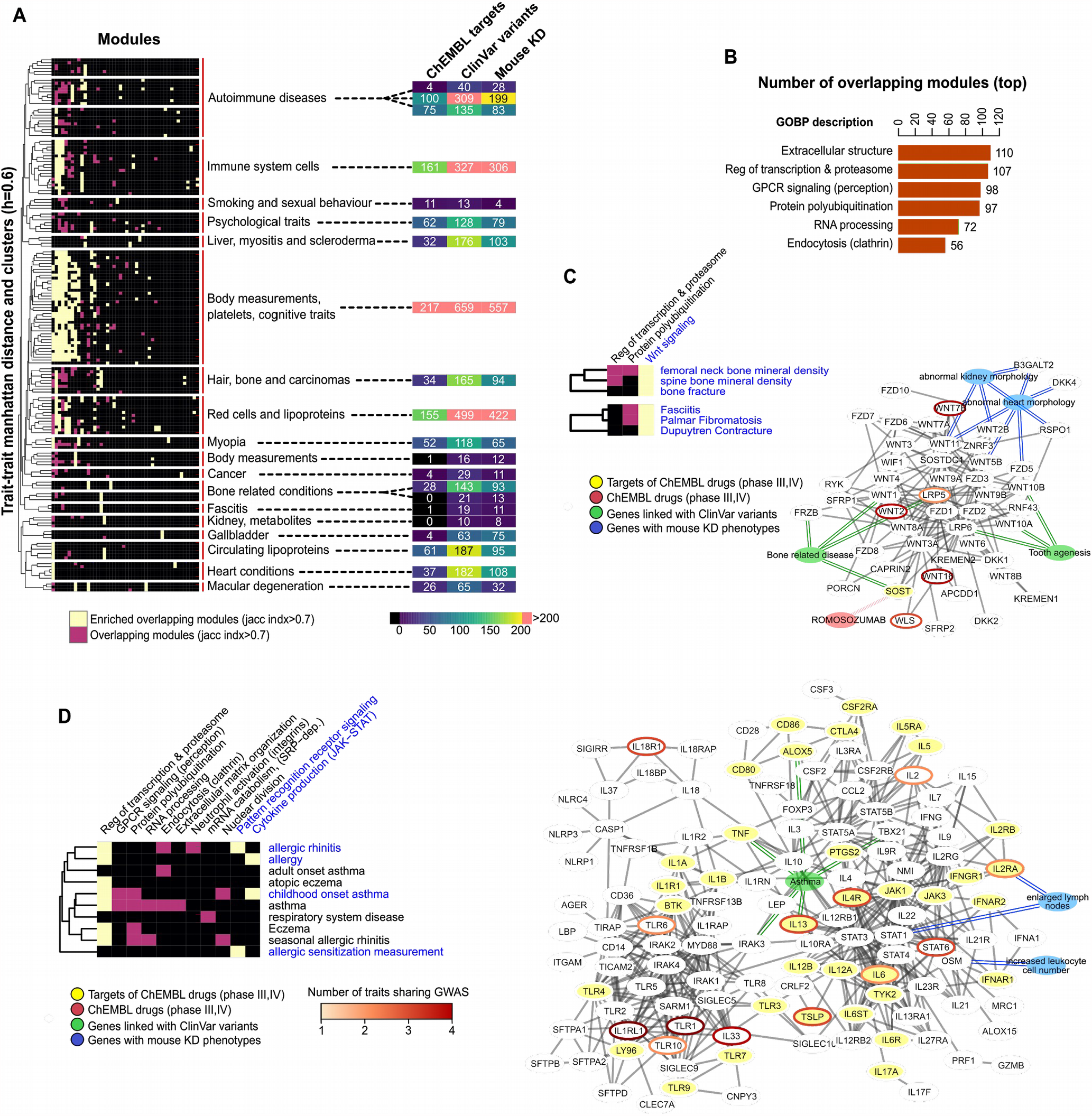
Multi-trait gene module associations for studies of shared biological processes and drug repurposing opportunities. A) heatmap showing the overlap between gene modules across traits. The traits were clustered by hierarchical clustering (see Methods) and subgroups defined by a cut-off of 0.6 average correlation coefficient. A module was considered the same across different traits when most genes are in common (Jaccard index > 0.7). Significant trait-module relations are marked in yellow or pink with yellow marking modules overrepresented in one of the sub-groups of traits (fisher test, adjusted p-value<0.05), and pink otherwise. The heatmap in the right panel shows the number of genes in modules from each sub-group of traits which are drug targets (phases III or higher, ChEMBL database), linked with clinical variants (ClinVar database), or with mouse knock-out phenotypes (IMPC database). B) Barplot showing the number of traits linked with the top six most pleiotropic gene modules. The Gene Ontology Biological process (GOBP) description is based on the results of a GOBP enrichment test (see Methods). C) Simplified heatmap of the clusters in figure A concerning bone related and fasciitis traits. The represented network includes genes from the modules indicated in blue letters and the represented interactions have been filtered for visualization (see Methods). Blue nodes - relevant mouse KO phenotypes; Green nodes - diseases with clinical variants enriched in this gene module; red nodes - drugs in clinical trials. Genes linked to blue, green or yellow nodes have the linked mouse phenotypes, clinical variants in the linked disease or are targets of the linked drug. Genes that are targets of drugs in clinical trials have yellow nodes. GWAS linked genes (L2G score >0.5) have borders coloured in an orange to red gradient (count of GWAS linked traits). D) Simplified heatmap of one the clusters in figure A concerning allergic reactions (the same node and edge color code as in C applies). In this case two modules were merged for building the interaction network in the right panel.

The six most pleiotropic gene modules were linked to between 56 and 110 traits in our study, and were enriched (GOBP enrichment with fisher test, BH adjusted *p-value* <0.05) for genes involved in protein ubiquitination, extracellular matrix organization, RNA processing and GPCR signalling (**Fig 3B**). These observations are in line with gene deletion studies in yeast that have identified some of the same cellular processes as highly pleiotropic (Hillenmeyer et al. 2008). Targeting pleiotropic processes with drugs could have broad applications but may also raise important safety concerns. To study this we obtained human genetic-interaction data (see Methods) and we observed that genes within the 73 gene modules linked to multiple traits have a small but significant increase in the average number of genetic interactions (enrichment of genetic interactions, Fisher test *p-value* = 4.155×10^−10^).

The traits linked with the 73 pleiotropic gene modules (shared between 2 or more traits) tend to have a higher number of significant initial GWAS seed genes (**SFig 1**). This difference is even more pronounced for the six most pleiotropic gene modules (**SFig 1**). Therefore, traits with a larger number of linked loci are more likely to be associated with pleiotropic gene modules. The 73 pleiotropic gene modules tend to be grouped according to coherent biological themes such as immune diseases, body measurements, and bone related conditions (**Fig 3A**). For each of these groups we then highlighted the gene modules that are over-represented in each group of traits (**Fig 3A**, Methods, fisher test, BH adjusted *p-value* < 0.05). To facilitate the study of cell biology and drug repurposing opportunities we have annotated (**Fig 3A**, and **STable 2**) the genes found in overlapping modules for each of the clusters with data from: ChEMBL (targets of drugs in at least phase III clinical trials), ClinVar (genes linked to clinical variants) and mouse knock-out phenotypes (phenotypic relevance and possible biological link). We explore a few examples of these modules in the following sections.

### Examples of shared molecular mechanisms and drug repurposing opportunities

We identified two groups of traits (bone and fasciitis related traits) which are predicted to have a common determining gene module (**Fig 3C** and **STable 3**). This module is enriched in Wnt signalling genes, which have been previously linked to bone homeostasis (Baron and Kneissel 2013) and to different types of fasciitis as well as Dupuytren’s contracture (Balaji, Kaveri, and Bayry 2011). We collected from ClinVar genes harbouring likely pathogenic variants found in patients (see Methods), hereafter referred to as ClinVar variants. This gene module is enriched in genes harbouring ClinVar variants from patients with tooth agenesis and bone related diseases (osteoporosis and osteopenia). Several genes with ClinVar variants associated with these diseases, such as *LRP6, SOST, WNT1, WNT10A* and *WNT10B*, are not linked to bone diseases via GWAS. Genetic manipulation of several genes within this module causes changes in bone density in mouse models (Wang et al. 2014). In addition, this module contains the target (SOST) of Romosozumab, a drug proven effective to treat osteoporosis. The ClinVar variants, mouse genetic models and the Romosozumab drug serve as an independent validation of the importance of this gene module for these traits. It also provides a proof of principle example of how this approach may be used to study the cell biology underlying a group of related traits and to identify relevant drug targets.

A second example (**Fig 3D** and **STable 3**) demonstrates the potential of multi-trait gene module associations for drug repurposing. We identified a group of ten respiratory (e.g. asthma) and cutaneous (e.g. eczema) immune-related diseases that share three gene modules - a highly pleiotropic module related to regulation of transcription and proteasome, and two more specific modules related to pattern recognition receptor signalling and cytokine production with JAK-STAT involvement. Genes in these modules had a significant enrichment (fisher test, *p-value* <0.05) in genes having likely pathogenic variants from patients with asthma. The two most specific gene modules were grouped together and shown in **Fig 3D** highlighting several genes with known pathogenic variants not associated with these diseases via GWAS (e.g. *IRAK3, TNF, ALOX5, TBX21*). *IRAK3*, encoding a protein pseudo-kinase, is an example of a druggable gene not identified by GWAS for asthma, but with protein missense variants linked to this disease (Balaci et al. 2007) and mice model studies implicating the regulation of IRAK3 in IL-33 induced airway inflammation (Nechama et al. 2018). While no drug for IRAK3 is used in the clinic, this analysis suggests it may serve as a relevant drug target for asthma and other related diseases.

We identified a total of 41 targets of 126 drugs targeting the genes in the module from **Fig 3D**. To identify drugs that could have repurposing potential, we excluded drugs already targeting therapeutic areas that include the 10 diseases linked to this gene module. This resulted in 18 drugs (**STable 3**) targeting 5 genes including: 14 drugs targeting *PTGS2*, used to treat primarily rheumatic disease and osteoarthritis; interferon alfacon1 or alfa-2B (targeting *IFNAR1* and *IFNAR2*), designed to counteract viral infections; galiximab and antibody for *CD80* (phase III trials for lymphoma); and the antibody RA-18C3 targeting IL1A for colorectal cancer. These drugs may be relevant to repurpose for respiratory or cutaneous autoimmune-related diseases. As a relevant example, RA-18C3 has shown benefit in a small phase II trial for hidradenitis suppurativa (acne inversa) (Gottlieb et al. 2020).

### Gene module analysis of genetically related immune-mediated diseases

Immune system related traits are well represented in our analysis, falling into three different groups: one containing systemic and organ-specific diseases, one cluster of immune cell measurements and a third more heterogeneous cluster (**Fig 3A, STable 2**). In **Fig 4A** we represent the first of these that can be further subdivided into: a sub-group linking the inflammatory bowel diseases (IBD), Multiple Sclerosis (MS) and Systemic Lupus Erythematosus (SLE); and subgroup linking celiac disease (CeD), Vitiligo and others. We find six gene modules that are specifically enriched with at least one of these two groups of traits, including gene modules related with GPCR signalling, neutrophil activation and interferon signalling. Genes present in these modules show higher relative expression (**Fig 4A**, right) in key immune tissues.

**Figure 4.**
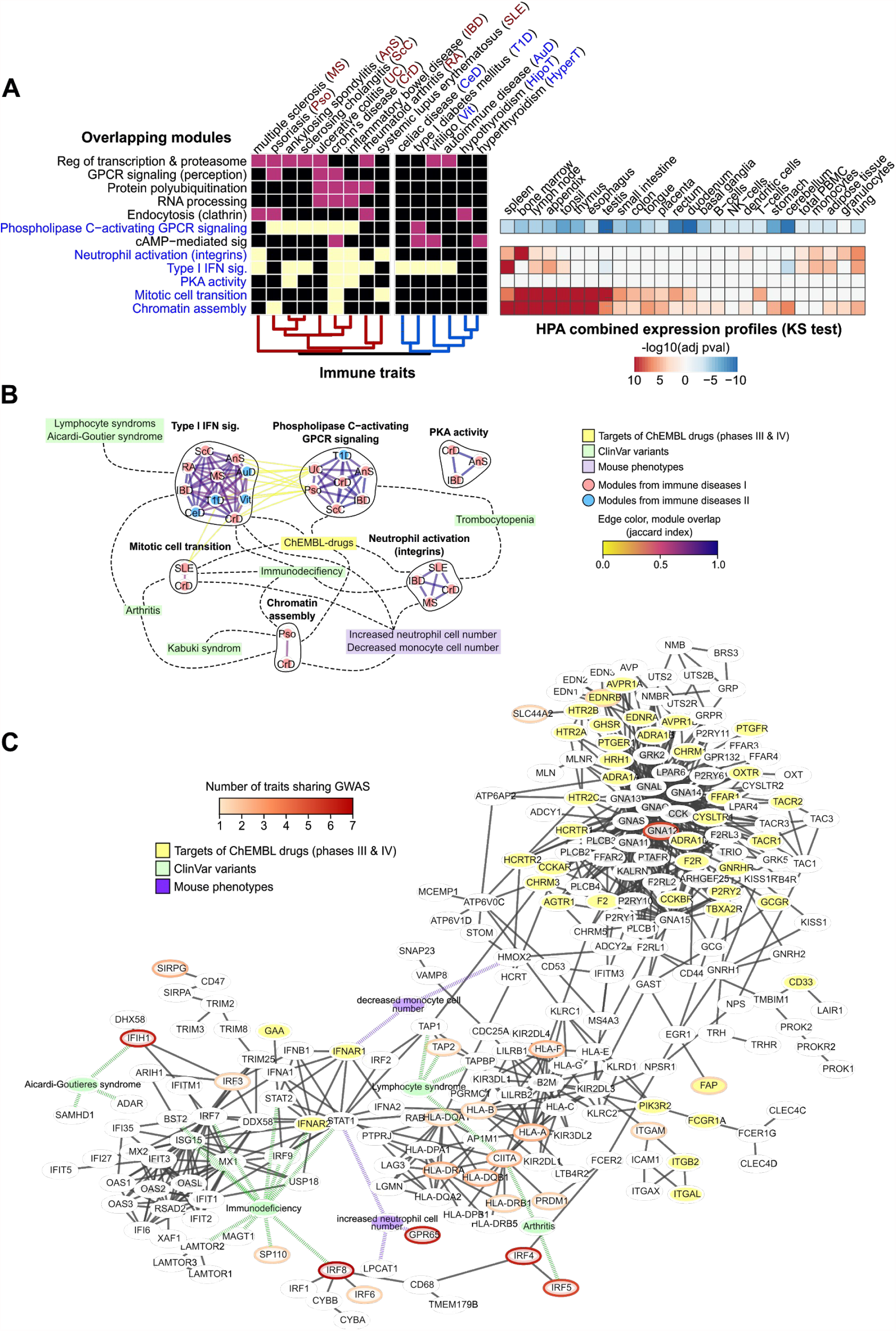
Gene module analysis of autoimmune diseases. A) Heatmap showing the overlap between gene modules across traits. The GOBP description is based on the results of a GOBP enrichment test (see material and methods). The heatmap in the right panel shows the gene set enrichment analysis done in the expression data from different tissues extracted from Human Protein Atlas for the gene modules in blue letters (see Methods). After BH adjustment for multiple testing, the p-value of the test was log transformed and given a positive value if the median distribution for the foreground is higher than the background and negative for the opposite. B) Shared modules as a network, nodes are gene modules associated with different immune related traits coloured in blue or red for the two trait sub-groups, edges represent high overlap at gene-level (Jaccard index>0.7). Gene modules linked to different traits are contained in black circles. Gene modules are linked with the yellow nodes “ChEMBL-drugs” when they contain targets for drugs in clinical trials (phases III and IV, ChEMBL); linked with green nodes when they are enriched in genes with clinical variants for a given disease; and linked to purple nodes when they are enriched for the corresponding KO phenotypes (fisher test, adjusted p-value<0.05). C) Network corresponding to genes found in gene modules enriched for Type I INF signalling, PLC activating GPCR signalling, Neutrophil activation (integrins) and PKA activity. Edge filtering, node and edge colours are the same as in figure 3 C-D.

To visualize the relationships between the traits and the 6 gene modules, we graphically linked the gene modules when there was a significant gene level overlap (**Fig 4B**, see Methods). Genes from these six modules showed enrichments for ClinVar variants from patients with immune diseases and relevant mouse gene KO phenotypes, further validating our approach. To represent the gene networks most relevant for these diseases we selected genes from modules linked with at least three immune-mediated diseases corresponding to those enriched in *Type I INF signalling, PLC activating GPCR signalling, Neutrophil activation (integrins)* and *PKA activity*. For representation (**Fig 4C**) we kept a subset of interactions of high confidence (see Methods) and highlighted genes with relevant ClinVar variants (green), mouse phenotypes (purple) and drug targets (yellow). We find multiple genes with ClinVar variants from patients with primary immune deficiencies (e.g. *IRF9, IRF7, STAT1, STAT2*) that are not GWAS linked genes but are in the network vicinity of those, providing further evidence of the importance of this gene module for these diseases.

To pinpoint drugs with repurposing potential, we excluded drugs targeting diseases in the same therapeutic areas shared by the immune mediated group of diseases, identifying 49 drugs with 20 targets. These include ulimorelin, an agonist of the ghrelin hormone secretagogue receptor *GHSR* used to treat gastrointestinal obstruction. Ghrelin hormone signalling has been studied in the context of age-related chronic inflammation (C. Fang et al. 2018), psoriasis (Qu et al. 2019) and IBD (reviewed in (Eissa and Ghia 2015)) indicating a potential repurposing opportunity. The 49 drugs with repurposing potential are listed in **STable 3** with information on target genes and clinical trials.

### Prioritization of IBD GWAS candidate genes using a network-based approach

Although the gene modules we have described can highlight biological pathways shared between genetically-related traits, identifying causal genes at individual GWAS loci is important for prioritising therapeutic targets. Existing methods such as GRAIL (Raychaudhuri et al. 2009), DEPICT (Pers et al. 2015), and MAGMA (de Leeuw et al. 2015) prioritise genes based on annotated or inferred biological pathways. However, they do not fully use the available genome-wide protein interaction networks, which can provide finer-grained resolution than gene sets grouped by gene ontology terms.

Here we use network propagation to prioritise genes at IBD GWAS loci, similar to the approach developed in our previous work on Alzheimer’s disease (Schwartzentruber et al. 2021). We used two alternative methods of defining seed genes for the network: first, we manually curated 37 genes with high confidence of being causally related to either Crohn’s disease or ulcerative colitis (Supplementary Table 4); second, we used the Open Targets L2G score to automatically select 110 genes with L2G > 0.5 at established IBD loci (Liu et al. 2015; de Lange et al. 2017) (see Methods; **STable 4**). To obtain network propagation scores unbiased by node degree, we compared each gene’s score to 1000 runs using the same number of randomly selected input genes, giving a Pagerank percentile value (see Methods). We obtained unbiased network propagation values for each seed gene by excluding each seed gene one at a time (see Methods).

We found that our curated seed genes had far higher network scores than other genes within 200 kb (p = 7.4×10^−6^, one-tailed Wilcoxon rank sum test), indicating that the majority of them have close interactions with other seed genes (**Fig 5A**). The same was true when considering seed genes exclusively in the L2G gene set (**Fig 5B**; *p-value*=3×10^−10^, one-tailed Wilcoxon rank sum test), indicating that many of these are also strong IBD candidate genes. Finally, we examined the enrichment of low SNP *p-values* within 10 kb of genes having high network scores. This revealed a progressive enrichment of low *p-values* near genes with higher network scores (**Fig 5C**), which held for the large number of genes linked to SNPs not reaching the typical genome-wide significance threshold of 5×10^−8^ for locus discovery.

**Figure 5.**
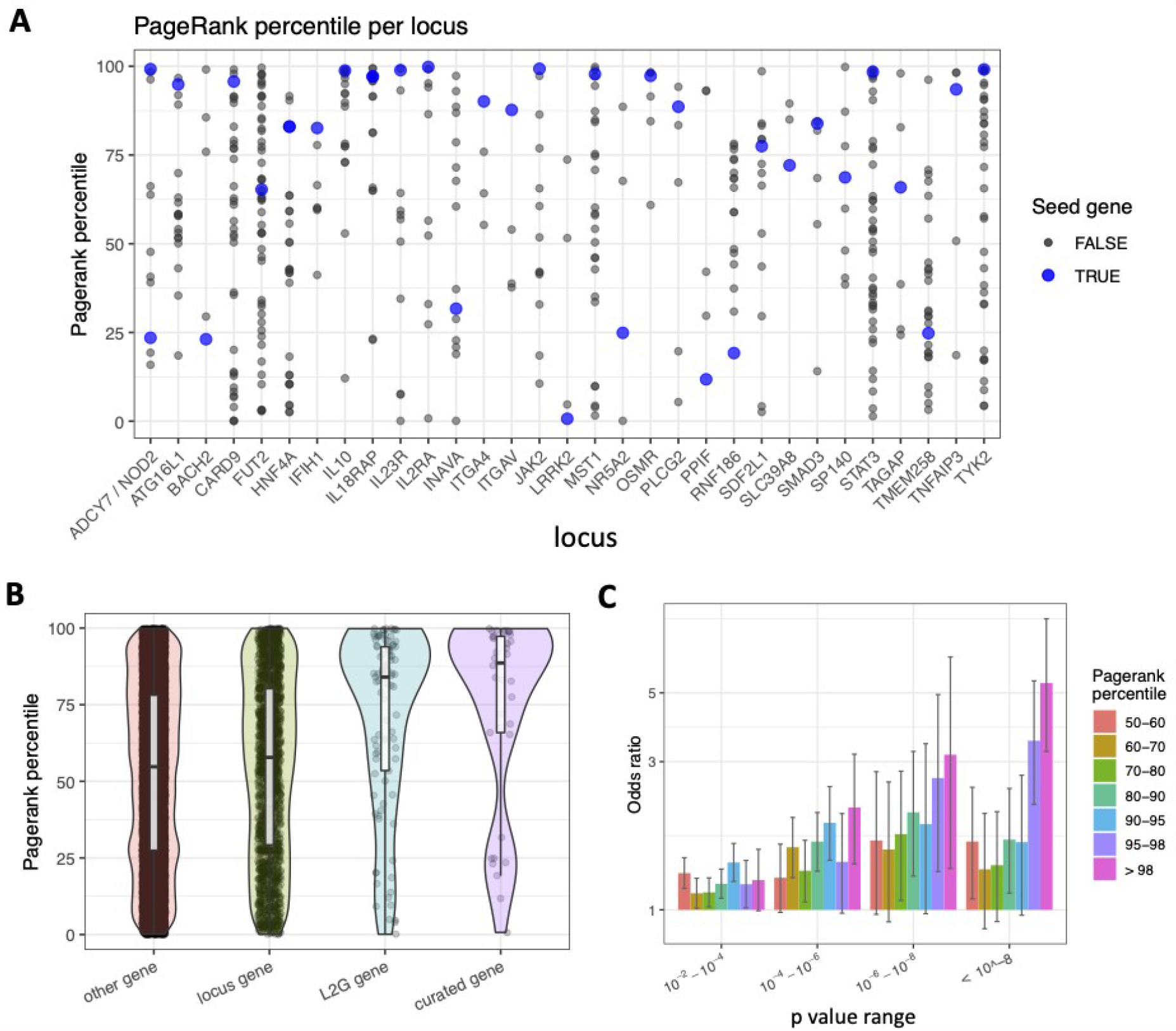
An IBD-specific network is enriched for likely causal genes. A) Curated IBD seed genes tend to have higher network propagation score (i.e. pagerank percentile) than other genes within 200 kb at the same loci. B) Genes selected by high Open Targets L2G score also tend to have high pagerank percentile, highlighting network evidence as complementary to typical locus features. C) Genome-wide, genes with low p-value SNPs within 10 kb are enriched for having high pagerank percentile.

Genes with the strongest network support included *TYK2* and *ICAM1*, both targets of drugs used for IBD (Tofacitinib and Natalizumab, respectively). Other curated IBD-causal genes with strong network support included *NOD2* and *IL23R*, which have missense variants implicating them as modulators of IBD (Hugot et al. 2001; Ogura et al. 2001; Duerr et al. 2006), and the drug target *ITGA4* (Vedolizumab). A small number of curated genes had lower network support. For example, *PPIF* encodes cyclophilin D, which regulates mitochondrial membrane potential. Recent CRISPRi-FlowFISH experiments showed an effect of the enhancer harboring IBD-associated variants specifically in stimulated immune cell types, providing a strong link with IBD pathogenesis. The lack of network support for *PPIF* could indicate that this gene affects IBD via pathways distinct from the biological functions most well covered by the curated gene set. Another curated IBD gene, *BACH2*, is associated with lymphoid cell counts and multiple autoimmune diseases (Vuckovic et al. 2020). *BACH2* receives a low score in the curated seed gene network, but a very high score in the L2G seed gene network, due to its interaction with candidate genes *FOSL2* and *PRDM1*. Overall there is moderate correlation between gene scores for the two sets of seed genes (spearman rho = 0.54), suggesting that it is useful to look within networks based on highly selected seed genes as well as network scores based on a broader set of candidate genes.

Across IBD loci without curated effector genes, our network scores provide further evidence to 42 candidates as being more highly functionally connected than remaining genes at the locus (**STable 4**, Methods). For each of the 42 genes we counted the number of traits with L2G score greater than 0.5 (**STable 4**) noting that 21 are linked to 10 or more traits and 12 to more than 20 different traits. While many of these were already strong IBD candidate genes that didn’t meet our top confidence threshold, some have only recently found strong support. A clear example is the *RIPK2* locus. Although *OSGIN2* is nearest to IBD lead SNP rs7015630 (38 kb distal), it has no apparent functional links with IBD (network score 43%). In contrast, *RIPK2* (108 kb distal, network score 99%) encodes for a mediator of inflammatory signalling via the interaction with the bacterial sensor *NOD2* (Canning et al. 2015). Network information can also provide a comparison point for other evidence sources. At the *DLD-SLC26A3* locus, there is moderate evidence of genetic colocalization between IBD and an eQTL for *DLD* in various tissues (Open Targets genetics portal). However, DLD has no clear functional links with IBD and receives a low network score (14%). In contrast, *SLC26A3* is a chloride anion transporter highly expressed in the human colon, with a high network score (98.4% in the L2G seed gene network), and its expression has been recently associated with clinical outcomes in ulcerative colitis (Camarillo et al. 2020). IBD candidate genes that have high network scores but have not been well characterized in the context of IBD include *PTPRC* (a phosphatase required for T-cell activation) and *BTBD8. BTBD8* is not a well studied gene (3 publications in Pubmed), but it is functionally connected to autophagy by the network analysis (via *WIPI2* and *ATG16L1*).

These results provide further evidence to candidate genes for IBD studies and drug development and illustrate the potential of integrating the network propagation scores as part of fine-mapping and gene prioritization efforts across other traits.

## Discussion

We identified gene modules associated with 906 human traits, taking advantage of the increase in coverage of human interactome mapping and novel tools for SNP to gene mapping (Mountjoy et al. 2020). As seen in other studies (Huang et al. 2018), network expansion is capable of retrieving previously known disease genes and drug targets that are not identified by GWAS. Network expansion can lead to the indication of genes that are not in GWAS loci but that may regulate or modulate the same biological processes. Importantly, even when excluding genes with direct genetic support, such interacting genes are enriched for successful drug targets (MacNamara et al. 2020). Genes identified by network expansion will not have information on direction of effect and additional work and interpretation is needed to gain insights into the direction of impact of modulating such genes.

Improvements in SNP to gene mapping provided a small but measurable improvement in the results of the network expansion when compared to using the closest gene to the identified SNP. While there are several algorithms to perform network propagation, recent studies have shown that they tend to perform similarly (Choobdar et al. 2019) and instead the network used has a stronger impact on performance (Huang et al. 2018). For this reason, improvements in mapping coverage and computational or experimental approaches to derive tissue or cell type specific networks (Greene et al. 2015) could have a large impact on future effectiveness of network expansion.

We showed examples of disease-linked gene modules that were also enriched in genes carrying clinical variants for the same or related diseases. In many cases, the genes with clinical variants did not overlap with the GWAS linked genes, which is likely due to lower frequency of clinical variants. Testing for burden of loss-of-function (LoF) variants within selected gene-sets is an approach used to study the impact of low frequency variants (Epi4K consortium and Epilepsy Phenome/Genome Project 2017; Povysil et al. 2019). We suggest that the gene modules identified here could be ideally suited for testing the burden of LoF in population scale genome sequencing efforts.

The gene modules identified here relate specific aspects of cell biology with different human traits. The analysis of mouse phenotypes and ClinVar variants provided additional evidence for some of the identified relationships. Additional work, in particular with appropriate models (e.g. organoids, mouse models) will be needed to follow up on some of the derived associations. The most pleiotropic gene modules reflect aspects of cell biology that have been defined as highly pleiotropic in gene deletion studies of yeast (Hillenmeyer et al. 2008). Interestingly, the traits that are linked with highly pleiotropic gene modules tend to have a larger number of starting GWAS seed genes. This suggests that the larger the number of loci linked to a trait the higher the chances that this trait will be genetically linked to a small number of highly pleiotropic biological processes. While it has been suggested that the heritability of complex traits is broadly spread along the genome (Boyle, Li, and Pritchard 2017), our analysis indicates that, across a large number of traits, this heritability overlaps in a non random fashion.

Gene modules linked with different traits could provide opportunities for drug repurposing or cross-disease drug development. However, pleiotropic effects of perturbing the related cell biological processes could raise safety concerns. Despite this, we did not find a strong correlation between the number of traits associated with a gene module and quantitative metrics relating to drug safety. Beyond identifying gene modules, our GWAS-based network approach can also be used to prioritise disease genes at individual loci by their role within specific biological processes, as we showed for IBD.

In summary, network expansion of GWAS is a powerful tool for the identification of genes and cellular processes linked to human traits, and the application to multi-trait analysis can reveal pleiotropy among human biological pathways, as well as highlight new opportunities for drug development and repurposing.

## Methods

### Human interactome, GWAS traits and linked genes analyzed

We created a comprehensive human interactome, merging an interactome developed for the Open Targets (www.opentargets.org) project (version from November 2019), with STRING v11.0. The Open Targets Interactome network was constructed during this project and contains human data only, including physical interaction data from IntAct, causality associations from SIGNOR and binarized pathway reaction relationships from Reactome. More details about the network construction can be found here: https://platform-docs.opentargets.org/target/molecular-interactions. STRING functional interactions were only human and selected to have a STRING edge score >=0.75. All identifiers were mapped to Ensembl gene identifiers and after removing duplicated edges and self-loops the final network used contains 18,410 nodes and 571,917 edges.

### Network propagation of GWAS linked genes

From a total number of 1221 traits, we selected 1,002 mapped to EFO terms (www.ebi.ac.uk/efo/) included in the Open Targets genetic portal, with at least 2 genes mapped to our interactome with a Locus to Gene score (L2G) of at least 0.5 (defined as seed nodes). The network-based approach was run individually for each trait, with each protein having a weight corresponding to the L2G score (between 0.5 and 1.0). The input was diffused through the interactome using the Personalized Page Rank algorithm (PPR) included in the R package igraph (v.1.2.4.2). To generate the modules, we selected the nodes with a PPR ranking score bigger than the third quartile (Q3, 75%) and performed walktrap clustering (igraph v.1.2.4.2). When the number of nodes in one module was bigger than 300, we repeated the clustering inside this community, until all resulting clusters were smaller than 300 genes. To define gene modules as significantly associated with a trait, we used a Kolmogorov Smirnov test to determine whether ranks (based on PPR) of genes in a module were greater than the background ranks of all the nodes considered for the walktrap clustering. We only tested modules with at least 10 genes and where at least 2 of them were seed genes (i.e. L2G>0.5), and we corrected the resulting *p-values* for multiple testing using BH adjustment. Based on this we identified a total of 2021 associations between a gene module and a trait.

### Benchmarking the capacity to predict disease associated genes from the network expansion

To benchmark both the predictive power of the ranking score resulting from the PPR and the genetic portal data when compared to GWAS catalog (https://www.ebi.ac.uk/gwas/, based on gene proximity), we computed ROC curves using as true positives the genes linked to diseases from the Jensen lab DISEASE database (diseases.jensenlab.org). This database provides a score measuring this association, the benchmark was done using 4 different score threshold (DIS0: all genes, DIS1: score>25%, DIS2: score>50%, DIS3: score>75% and DIS4: maximum value for the score). We calculated the ROC curves and the AUCs (area under the ROC curve) for traits with at least 10 True Positives. Also, we randomized both the nodes in the network (keeping the degree distribution) as well as the true positives 1,000 times each, then we calculated the AUCs and the subsequent Z Scores. As an extra benchmark we used the clinical trial data contained in ChEMBL, considering as true positives drug targets tested for a certain disease at clinical phases II or higher.

### Trait-trait relationships defined by the similarity of the network propagation

We calculated the Manhattan distance between the 1002 traits using the full PPR ranking score, followed by hierarchical clustering, resulting in 54 clusters (height distance=1). To further characterize them, we selected the ones having at least 5 traits, we obtained their EFO ancestry and calculated their frequency per cluster. The highest frequency per cluster is used to define 9 groups colour coded in Fig 2A. To complement the description of clusters belonging to the most general group “measurement” and “material property”, we extracted EFO ancestry terms with manually assigned terms from the EFO ancestry with lower frequency and listed in Fig 2A. ChEMBL database was used to calculate the counts of both drugs and drug targets for each of the trait clusters, using the information for drugs in clinical trials, phases III and IV. To further illustrate the validity of this approach, we selected 3 trait clusters (Fig 2B) as examples of valid trait to trait relations.

### Multi-trait gene module analysis

The significant modules identified for each trait (described above) were compared across traits by measuring the overlap in genes using the Jaccard index. Gene modules with Jaccard index >= 0.70 were considered to be in common across two traits. From the 2021 pairs of gene modules to trait associations, 886 are unique to a single trait and the remainder can be collapsed to 73 gene modules that are enriched in network propagation signals for 2 or more traits. To identify which sub-groups of related traits we clustered the traits linked to the 73 multi-trait modules based on the Manhattan distance of their full PPR ranking score (as above) using hierarchical clustering. Sub-groups were defined with a height cut-off of 0.7 and we identified gene modules that were more specific to each sub-group of traits using a fisher test and BH multiple testing correction. We kept trait sub-groups with at least 3 traits and significant presence of at least one group of overlapping modules.

### Gene module annotations and enrichment analysis

The gene KD mouse phenotypes were extracted from the International Mouse Phenotyping Consortium (IMPC) and the clinical variants from the database ClinVar (NCBI). For the enrichment of genes from clinical variants, the diseases were grouped into larger categories. For the enrichment of genes from clinical variants referred to in Fig 3C-D and Fig 4 B-C, we downloaded the data from ClinVar (NCBI), filtered out all benign associations and grouped the phenotypes larger higher categories as follows: tooth agenesis (tooth agenesis, Selective tooth agenesis 4, 7 and 8), bone related diseases (sclerosteosis 1, osteoarthritis, osteopetrosis, osteoporosis, osteogenesis imperfecta and osteopenia), asthma (asthma and nasal polyps, susceptibility to asthma and asthma related traits, diminish response to leukotriene treatment in asthma, asthma and aspirine intolerance), autoimmune condition (Familial cold autoinflammatory syndromes), immunodeficiency (immunodeficiency due to defect in mapbp-interacting protein, hepatic venoocclusive disease with immunodeficiency, immunodeficiency-centromeric instability-facial anomalies syndrome 1, immunodeficiency 31a, 31C, 32a, 32b, 38, 39, 44 and 45, immunodeficiency X-Linked, with magnesium defect, Epstein-Barr virus infection, and neoplasia, combined immunodeficiency, severe T-cell immunodeficiency, and immunodeficiency 65 with susceptibility to viral infections), lymphocyte syndrome (Bare lymphocyte syndrome types 1 and 2), arthritis (rheumatoid arthritis and juvenile arthritis), Kabuki syndrome (Kabuki syndrome 1 and 2), thrombocytopenia (thrombocytopenia, dyserythropoietic anaemia with thrombocytopenia, GATA-1-related thrombocytopenia with dyserythropoiesis, X-linked thrombocytopenia without dyserythropoietic anaemia, thrombocytopenia with platelet dysfunction, hemolysis, and imbalanced globin synthesis, radioulnar synostosis with amegakaryocytic thrombocytopenia 2 and macrothrombocytopenia), anaemia (anaemia, dyserythropoietic anaemia with thrombocytopenia, aplastic anaemia, CD59-mediated haemolytic anaemia with or without immune-mediated polyneuropathy and Diamond-Blackfan anaemia) and aicardi-Goutieres syndrome (Aicardi-Goutieres syndrome 4, 6 and 7).

### IBD network analyses for fine-mapping

To identify robust IBD-associated loci, we extracted loci defined in the Open Targets Genetics portal (genetics.opentargets.org) for two IBD GWAS (de Lange et al. 2017; Liu et al. 2015). Since each GWAS may identify different lead variants, we merged together loci defined by lead variants within 200 kb of each other. We extracted the locus2gene (L2G) score reported for all genes at each locus, and for merged loci took the average L2G score for each gene across the loci. We curated 37 high-confidence IBD genes based on the presence of fine-mapped deleterious coding variants, genes whose protein products are the targets of approved IBD drugs, and literature. We defined additional seed gene sets by selecting the top gene at each locus which had a L2G score > 0.5. We ran network propagation as described in the main text. However, to get unbiased scores for seed genes themselves, we left each seed gene out of the input in turn, and ran network propagation to obtain a score based on the remaining N-1 seed genes. To compute the PPR percentile for seed genes, we used the PPR percentile from the single network propagation run where that seed gene was excluded from the input. For all other genes, we used the median PPR percentile across the N seed gene runs. Plots in Fig 5 are based on PPR percentiles from the curated seed gene network. To assess enrichment of low *p-value* SNPs near high-network genes (Fig 5C), we first determined for each gene the minimum *p-value* among SNPs within 10 kb of the gene’s footprint based on IBD GWAS summary statistics from de Lange et al. (2017). We used Fisher’s exact test to determine the odds ratio for genes with high network score (in each defined bin) to have a low minimum SNP *p-value*, relative to genes with low network scores (PPR percentile < 50).

PPR percentiles discussed in the text are the average PPR percentiles for each gene across the curated and L2G>0.5 networks. We identified IBD candidate genes that stand out based on their network score (STable 4) by filtering all locus genes to those which had average PPR percentile > 90 and L2G > 0.1, and where no other gene at the same locus had PPR percentile > 80 and L2G > 0.1.

## Supplementary figures

**Supplementary figure 1.**
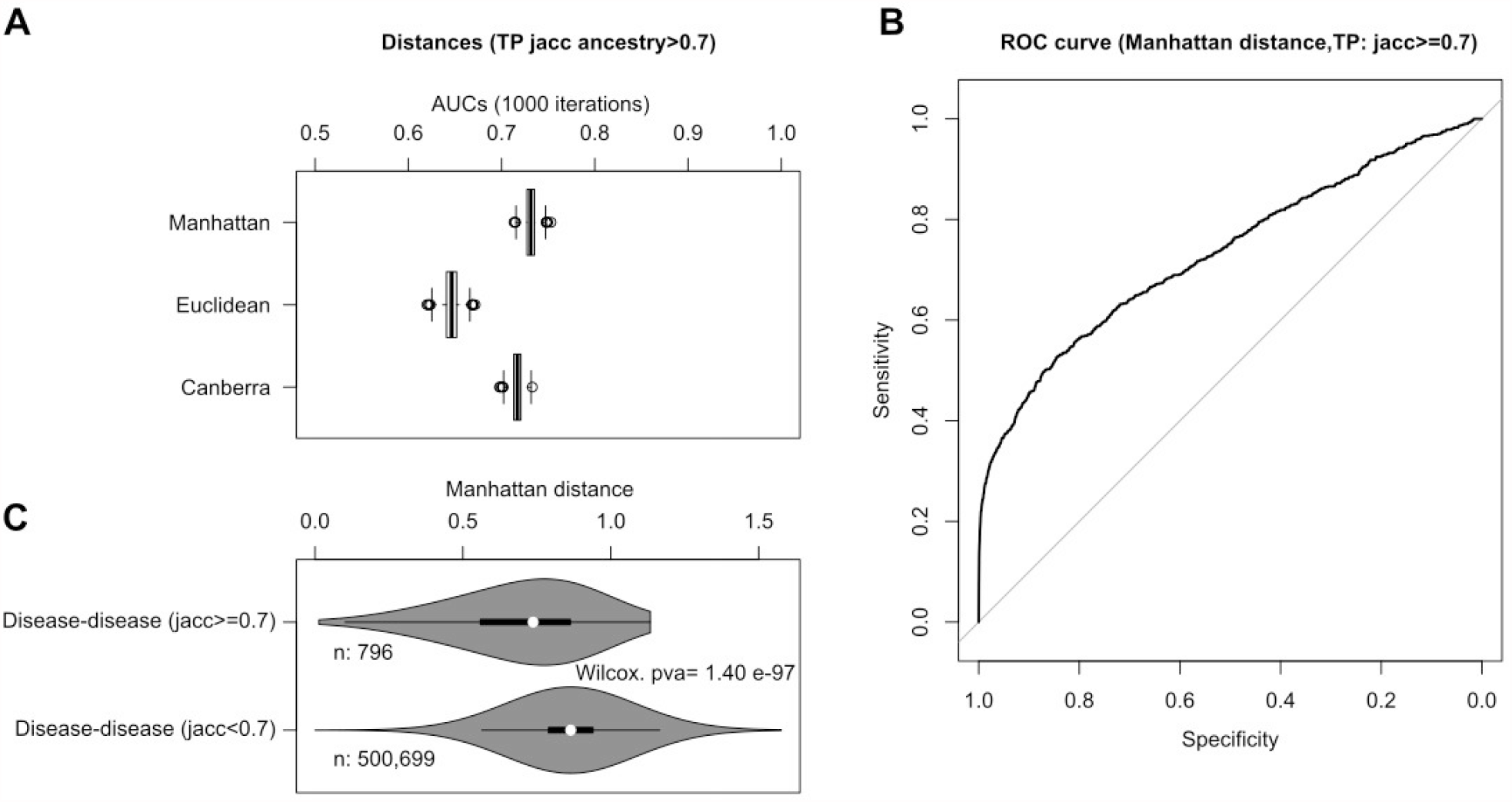
Disease-disease distance benchmark. A) Areas under the ROC curve (AUCs) for three different disease-disease distance metrics: Manhattan, Euclidean and Canberra distances. They were calculated using the full PPR ranking scores after the network expansion for all disease-disease pairs, we considered as true positives the 796 disease-disease pairs with common ancestry (Jaccard score of ancestry terms from EFO annotation bigger or equal to 0.7). To calculate the ROC curves, we sampled 1000 pairs from the negative space for 1000 iterations, the resulting AUCs were plotted in the boxplots. B) Example of one of the ROC curves for Manhattan distance (AUC= 0.73) C) Violin plot showing the Manhattan distance distribution for all disease-disease pairs with shared ancestry (jaccard index >=0.70) considered as true positive and for all pairs considered as negative space. The Wilcoxon rank sum test was calculated to measure the difference between both distributions.

**Supplementary figure 2.**
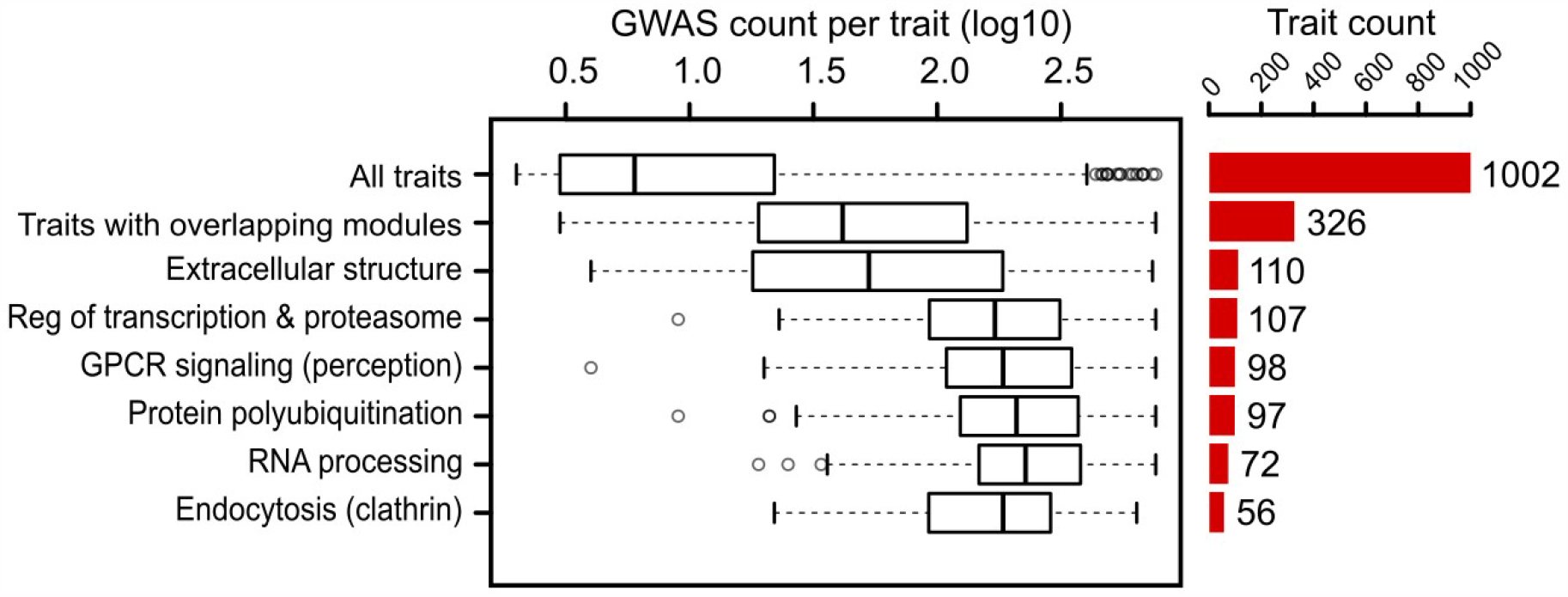
Boxplot showing the number of starting GWAS hits per trait (GWAS count) for all traits, for traits with shared modules and for traits that have highly pleiotropic modules (top 6, description based on GOBP annotation). In the left panel, barplot showing the total number of traits for each selection.

## Acknowledgements

JS, QZ and CAA are supported by Wellcome Trust Grant 206194. A CC BY or equivalent licence is applied to the AAM arising from this submission, in accordance with the grant’s open access conditions. IB is supported by funding from Open Targets.

## References

Balaci, Lenuta, Maria Cristina Spada, Nazario Olla, Gabriella Sole, Laura Loddo, Francesca Anedda, Silvia Naitza, et al. 2007. “IRAK-M Is Involved in the Pathogenesis of Early-Onset Persistent Asthma.” American Journal of Human Genetics 80 (6): 1103–14.

Balaji, Kithiganahalli N., Srini V. Kaveri, and Jagadeesh Bayry. 2011. “Wnt Signaling and Dupuytren’s Disease.” The New England Journal of Medicine.

Baron, Roland, and Michaela Kneissel. 2013. “WNT Signaling in Bone Homeostasis and Disease: From Human Mutations to Treatments.” Nature Medicine 19 (2): 179–92.

Boyle, Evan A., Yang I. Li, and Jonathan K. Pritchard. 2017. “An Expanded View of Complex Traits: From Polygenic to Omnigenic.” Cell 169 (7): 1177–86.

Camarillo, Gabriela Fonseca, Emilio Iturriaga Goyon, Rafael Barreto Zuñiga, Lucero Adriana Salazar Salas, Ana Elena Peredo Escárcega, and Jesús K. Yamamoto-Furusho. 2020. “Gene Expression Profiling of Mediators Associated with the Inflammatory Pathways in the Intestinal Tissue from Patients with Ulcerative Colitis.” Mediators of Inflammation 2020 (January): 9238970.

Canning, Peter, Qui Ruan, Tobias Schwerd, Matous Hrdinka, Jenny L. Maki, Danish Saleh, Chalada Suebsuwong, et al. 2015. “Inflammatory Signaling by NOD-RIPK2 Is Inhibited by Clinically Relevant Type II Kinase Inhibitors.” Chemistry & Biology 22 (9): 1174–84.

Carter, Hannah, Matan Hofree, and Trey Ideker. 2013. “Genotype to Phenotype via Network Analysis.” Current Opinion in Genetics & Development 23 (6): 611–21.

Choobdar, Sarvenaz, Mehmet E. Ahsen, Jake Crawford, Mattia Tomasoni, Tao Fang, David Lamparter, Junyuan Lin, et al. 2019. “Assessment of Network Module Identification across Complex Diseases.” Nature Methods 16 (9): 843–52.

Duerr, Richard H., Kent D. Taylor, Steven R. Brant, John D. Rioux, Mark S. Silverberg, Mark J. Daly, A. Hillary Steinhart, et al. 2006. “A Genome-Wide Association Study Identifies IL23R as an Inflammatory Bowel Disease Gene.” Science 314 (5804): 1461–63.

Eissa, N., and J. E. Ghia. 2015. “Immunomodulatory Effect of Ghrelin in the Intestinal Mucosa.” Neurogastroenterology and Motility: The Official Journal of the European Gastrointestinal Motility Society 27 (11): 1519–27.

Epi4K consortium, and Epilepsy Phenome/Genome Project. 2017. “Ultra-Rare Genetic Variation in Common Epilepsies: A Case-Control Sequencing Study.” Lancet Neurology 16 (2): 135–43.

Fang, Chuo, Hang Xu, Shaodong Guo, Susanne U. Mertens-Talcott, and Yuxiang Sun. 2018. “Ghrelin Signaling in Immunometabolism and Inflamm-Aging.” Advances in Experimental Medicine and Biology 1090: 165–82.

Fang, Hai, ULTRA-DD Consortium, Hans De Wolf, Bogdan Knezevic, Katie L. Burnham, Julie Osgood, Anna Sanniti, et al. 2019. “A Genetics-Led Approach Defines the Drug Target Landscape of 30 Immune-Related Traits.” Nature Genetics 51 (7): 1082–91.

Franke, Lude, Harm van Bakel, Like Fokkens, Edwin D. de Jong, Michael Egmont-Petersen, and Cisca Wijmenga. 2006. “Reconstruction of a Functional Human Gene Network, with an Application for Prioritizing Positional Candidate Genes.” American Journal of Human Genetics 78 (6): 1011–25.

Gottlieb, Alice, Nicola E. Natsis, Francisco Kerdel, Seth Forman, Edgar Gonzalez, Gilberto Jimenez, Liliam Hernandez, et al. 2020. “A Phase II Open-Label Study of Bermekimab in Patients with Hidradenitis Suppurativa Shows Resolution of Inflammatory Lesions and Pain.” The Journal of Investigative Dermatology 140 (8): 1538–45.e2.

Greene, Casey S., Arjun Krishnan, Aaron K. Wong, Emanuela Ricciotti, Rene A. Zelaya, Daniel S. Himmelstein, Ran Zhang, et al. 2015. “Understanding Multicellular Function and Disease with Human Tissue-Specific Networks.” Nature Genetics 47 (6): 569–76.

Hackinger, Sophie, and Eleftheria Zeggini. 2017. “Statistical Methods to Detect Pleiotropy in Human Complex Traits.” Open Biology 7 (11). https://doi.org/10.1098/rsob.170125.

Hillenmeyer, Maureen E., Eula Fung, Jan Wildenhain, Sarah E. Pierce, Shawn Hoon, William Lee, Michael Proctor, et al. 2008. “The Chemical Genomic Portrait of Yeast: Uncovering a Phenotype for All Genes.” Science 320 (5874): 362–65.

Huang, Justin K., Daniel E. Carlin, Michael Ku Yu, Wei Zhang, Jason F. Kreisberg, Pablo Tamayo, and Trey Ideker. 2018. “Systematic Evaluation of Molecular Networks for Discovery of Disease Genes.” Cell Systems 6 (4): 484–95.e5.

Hugot, J. P., M. Chamaillard, H. Zouali, S. Lesage, J. P. Cézard, J. Belaiche, S. Almer, et al. 2001. “Association of NOD2 Leucine-Rich Repeat Variants with Susceptibility to Crohn’s Disease.” Nature 411 (6837): 599–603.

Jassal, Bijay, Lisa Matthews, Guilherme Viteri, Chuqiao Gong, Pascual Lorente, Antonio Fabregat, Konstantinos Sidiropoulos, et al. 2020. “The Reactome Pathway Knowledgebase.” Nucleic Acids Research 48 (D1): D498–503.

Lange, Katrina M. de, Loukas Moutsianas, James C. Lee, Christopher A. Lamb, Yang Luo, Nicholas A. Kennedy, Luke Jostins, et al. 2017. “Genome-Wide Association Study Implicates Immune Activation of Multiple Integrin Genes in Inflammatory Bowel Disease.” Nature Genetics 49 (2): 256–61.

Lee, Insuk, U. Martin Blom, Peggy I. Wang, Jung Eun Shim, and Edward M. Marcotte. 2011. “Prioritizing Candidate Disease Genes by Network-Based Boosting of Genome-Wide Association Data.” Genome Research 21 (7): 1109–21.

Leeuw, Christiaan A. de, Joris M. Mooij, Tom Heskes, and Danielle Posthuma. 2015. “MAGMA: Generalized Gene-Set Analysis of GWAS Data.” PLoS Computational Biology 11 (4): e1004219.

Licata, Luana, Prisca Lo Surdo, Marta Iannuccelli, Alessandro Palma, Elisa Micarelli, Livia Perfetto, Daniele Peluso, Alberto Calderone, Luisa Castagnoli, and Gianni Cesareni. 2020. “SIGNOR 2.0, the SIGnaling Network Open Resource 2.0: 2019 Update.” Nucleic Acids Research 48 (D1): D504–10.

Liu, Jimmy Z., Suzanne van Sommeren, Hailiang Huang, Siew C. Ng, Rudi Alberts, Atsushi Takahashi, Stephan Ripke, et al. 2015. “Association Analyses Identify 38 Susceptibility Loci for Inflammatory Bowel Disease and Highlight Shared Genetic Risk across Populations.” Nature Genetics 47 (9): 979–86.

MacNamara, Aidan, Nikolina Nakic, Ali Amin Al Olama, Cong Guo, Karsten B. Sieber, Mark R. Hurle, and Alex Gutteridge. 2020. “Network and Pathway Expansion of Genetic Disease Associations Identifies Successful Drug Targets.” Scientific Reports 10 (1): 20970.

Mountjoy, Edward, Ellen M. Schmidt, Miguel Carmona, Gareth Peat, Alfredo Miranda, Luca Fumis, James Hayhurst, et al. 2020. “Open Targets Genetics: An Open Approach to Systematically Prioritize Causal Variants and Genes at All Published Human GWAS Trait-Associated Loci.” Cold Spring Harbor Laboratory. https://doi.org/10.1101/2020.09.16.299271.

Nechama, Morris, Jeahoo Kwon, Shuo Wei, Adrian Tun Kyi, Robert S. Welner, Iddo Z. Ben-Dov, Mohamed S. Arredouani, et al. 2018. “The IL-33-PIN1-IRAK-M Axis Is Critical for Type 2 Immunity in IL-33-Induced Allergic Airway Inflammation.” Nature Communications 9 (1): 1603.

Nelson, Matthew R., Hannah Tipney, Jeffery L. Painter, Judong Shen, Paola Nicoletti, Yufeng Shen, Aris Floratos, et al. 2015. “The Support of Human Genetic Evidence for Approved Drug Indications.” Nature Genetics 47 (8): 856–60.

Ogura, Y., D. K. Bonen, N. Inohara, D. L. Nicolae, F. F. Chen, R. Ramos, H. Britton, et al. 2001. “A Frameshift Mutation in NOD2 Associated with Susceptibility to Crohn’s Disease.” Nature 411 (6837): 603–6.

Orchard, Sandra, Mais Ammari, Bruno Aranda, Lionel Breuza, Leonardo Briganti, Fiona Broackes-Carter, Nancy H. Campbell, et al. 2014. “The MIntAct project—IntAct as a Common Curation Platform for 11 Molecular Interaction Databases.” Nucleic Acids Research. https://doi.org/10.1093/nar/gkt1115.

Oti, M., and H. G. Brunner. 2007. “The Modular Nature of Genetic Diseases.” Clinical Genetics 71 (1): 1–11.

Oti, M., B. Snel, M. A. Huynen, and H. G. Brunner. 2006. “Predicting Disease Genes Using Protein-Protein Interactions.” Journal of Medical Genetics 43 (8): 691–98.

Pers, Tune H., Genetic Investigation of ANthropometric Traits (GIANT) Consortium, Juha M. Karjalainen, Yingleong Chan, Harm-Jan Westra, Andrew R. Wood, Jian Yang, et al. 2015. “Biological Interpretation of Genome-Wide Association Studies Using Predicted Gene Functions.” Nature Communications. https://doi.org/10.1038/ncomms6890.

Porras, Pablo, Elisabet Barrera, Alan Bridge, Noemi Del-Toro, Gianni Cesareni, Margaret Duesbury, Henning Hermjakob, et al. 2020. “Towards a Unified Open Access Dataset of Molecular Interactions.” Nature Communications 11 (1): 6144.

Povysil, Gundula, Slavé Petrovski, Joseph Hostyk, Vimla Aggarwal, Andrew S. Allen, and David B. Goldstein. 2019. “Rare-Variant Collapsing Analyses for Complex Traits: Guidelines and Applications.” Nature Reviews. Genetics, October. https://doi.org/10.1038/s41576-019-0177-4.

Qu, Ruize, Xiaomin Chen, Jing Hu, Yufeng Fu, Jiangfan Peng, Yuhua Li, Jingxi Chen, et al. 2019. “Ghrelin Protects against Contact Dermatitis and Psoriasiform Skin Inflammation by Antagonizing TNF-α/NF-κBB Signaling Pathways.” Scientific Reports 9 (1): 1348.

Raychaudhuri, Soumya, Robert M. Plenge, Elizabeth J. Rossin, Aylwin C. Y. Ng, International Schizophrenia Consortium, Shaun M. Purcell, Pamela Sklar, et al. 2009. “Identifying Relationships among Genomic Disease Regions: Predicting Genes at Pathogenic SNP Associations and Rare Deletions.” PLoS Genetics 5 (6): e1000534.

Schwartzentruber, Jeremy, Sarah Cooper, Jimmy Z. Liu, Inigo Barrio-Hernandez, Erica Bello, Natsuhiko Kumasaka, Adam M. H. Young, et al. 2021. “Genome-Wide Meta-Analysis, Fine-Mapping and Integrative Prioritization Implicate New Alzheimer’s Disease Risk Genes.” Nature Genetics. https://doi.org/10.1038/s41588-020-00776-w.

Stacey, David, Eric B. Fauman, Daniel Ziemek, Benjamin B. Sun, Eric L. Harshfield, Angela M. Wood, Adam S. Butterworth, Karsten Suhre, and Dirk S. Paul. 2019. “ProGeM: A Framework for the Prioritization of Candidate Causal Genes at Molecular Quantitative Trait Loci.” Nucleic Acids Research 47 (1): e3.

Sun, Benjamin B., Joseph C. Maranville, James E. Peters, David Stacey, James R. Staley, James Blackshaw, Stephen Burgess, et al. 2018. “Genomic Atlas of the Human Plasma Proteome.” Nature 558 (7708): 73–79.

Szklarczyk, Damian, Annika L. Gable, David Lyon, Alexander Junge, Stefan Wyder, Jaime Huerta-Cepas, Milan Simonovic, et al. 2019. “STRING v11: Protein-Protein Association Networks with Increased Coverage, Supporting Functional Discovery in Genome-Wide Experimental Datasets.” Nucleic Acids Research 47 (D1): D607–13.

Vanunu, Oron, Oded Magger, Eytan Ruppin, Tomer Shlomi, and Roded Sharan. 2010. “Associating Genes and Protein Complexes with Disease via Network Propagation.” PLoS Computational Biology 6 (1): e1000641.

Vuckovic, Dragana, Erik L. Bao, Parsa Akbari, Caleb A. Lareau, Abdou Mousas, Tao Jiang, Ming-Huei Chen, et al. 2020. “The Polygenic and Monogenic Basis of Blood Traits and Diseases.” Cell 182 (5): 1214–31.e11.

Wang, Yiping, Yi-Ping Li, Christie Paulson, Jian-Zhong Shao, Xiaoling Zhang, Mengrui Wu, and Wei Chen. 2014. “Wnt and the Wnt Signaling Pathway in Bone Development and Disease.” Frontiers in Bioscience 19 (January): 379–407.

Watanabe, Kyoko, Sven Stringer, Oleksandr Frei Maša Umicevic Mirkov, Christiaan de Leeuw, Tinca J. C. Polderman, Sophie van der Sluis, Ole A. Andreassen, Benjamin M. Neale, and Danielle Posthuma. 2019. “A Global Overview of Pleiotropy and Genetic Architecture in Complex Traits.” Nature Genetics 51 (9): 1339–48.

Zhu, Zhihong, Futao Zhang, Han Hu, Andrew Bakshi, Matthew R. Robinson, Joseph E. Powell, Grant W. Montgomery, et al. 2016. “Integration of Summary Data from GWAS and eQTL Studies Predicts Complex Trait Gene Targets.” Nature Genetics 48 (5): 481– 87.

